# Emergence and persistence of the Chikungunya virus East-Central-South-African genotype in Northeast Brazil

**DOI:** 10.1101/111948

**Authors:** Antonio Charlys da Costa, Julien Thézé, Shirley Cavalcante Vasconcelos Komninakis, Rodrigo Lopes Sanz-Duro, Marta Rejane Lemos Felinto, Lúcia Cristina Corrêa Moura, Ivoneide Moreira de Oliveira Barroso, Lucineide Eliziario Correia Santos, Mardjane Alves de Lemos Nunes, Adriana Avila Moura, José Lourenço, Xutao Deng, Eric L. Delwart, Maria Raquel dos Anjos Silva Guimarães, Oliver G. Pybus, Ester C. Sabino, Nuno R. Faria

## Abstract

We investigate an outbreak of exanthematous illness in Maceió, Alagoas, using molecular surveillance. Of 273 samples, 76% tested RT-qPCR positive for Chikungunya virus. Phylogenetic analysis reveals that the outbreak was caused by the East-Central-South-African genotype, and that this lineage has likely persisted since mid-2014 in Northeast Brazil.

**Article summary line:** Transmission of the Chikungunya virus East-Central-South-African genotype has been ongoing in the Northeast region of Brazil since mid-2014.

## Article

Chikungunya virus (CHIKV) was first detected in the Americas in December 2013 (1). Transmission of this alphavirus (family *Togaviridae)* is mediated by infected anthropophilic vectors, such as *Aedes aegypti* and *Ae. albopictus.* CHIKV infection causes fever, rash and arthralgia; headache, conjunctivitis and joint swelling are also common. Complications of infection include Guillain-Barré syndrome and polyarthralgia that can last for months. The clinical symptoms of CHIKV infection are similar to those of dengue (DENV) and Zika viruses (ZIKV), complicating diagnosis in regions where these arboviruses co-circulate.

Between 30 March and 3 May 2016, a molecular investigation was conducted following an increase in emergency consultations due to exanthematous illness at two private hospitals in Maceió, capital city of Alagoas State, Brazil. Samples were obtained from 273 patients with on average 37 years old (from 1 to 86 years old); 64% (n=175/273) were female and 73% (n=198/273) of patients resided in Maceió municipality. The study was approved by the Faculdade de Medicina da Universidade de São Paulo Review Board, and informed consent was obtained from all subjects.

To avoid misdiagnosis based on clinical symptoms only (2), viral RNA was extracted from 140 μL of serum samples using QIAmp viral RNA kit (Qiagen) and RT-qPCR analyses were performed for DENV serotypes 1-4 (3), ZIKV (4) and CHIKV (5). Two additional CHIKV RT-qPCR+ samples from Paraíba were included in subsequent analysis. To identify the cause of the outbreak, each plasma sample was subjected to centrifugation at 15,000 x g for 10 minutes, filtered through a 0.45 μm filter (Millipore), then subjected to nuclease treatment, viral RNA extraction (Maxwell) and cDNA synthesis (Promega). A single round of DNA synthesis was performed using DNA Polymerase I Large (Klenow) Fragment (Promega). Subsequently, a Nextera XT Sample Preparation Kit (Illumina) was used construct a DNA library, with each sample identifiable using dual barcodes. The library was deep-sequenced using the MiSeq Sequencer (Illumina) with 300 base paired ends. BLASTx was used to identify viral sequences through their protein sequence similarity to annotated viral proteins in GenBank search, as described previously (6).

Of 273 samples tested, 76% (n=208) were CHIKV-RNA+ and 24% (n=66) were ZIKV-RNA+. Also, 13.2% of sampled (36/273) were co-infected of CHIKV and ZIKV, similar to recent findings from Salvador, Bahia (**7**). No DENV cases were detected in this population. Ct-values of CHIKV RNA+ samples were lower (n=208, average Ct-value=24.6) than those for ZIKV (n=66, average Ct-value=33.5).

A genetic analysis of all publicly available CHIKV genomes >1500nt in length (n=659 as of 17 Feb 2017) reveals that the East-Central-South-Africa (ECSA) genotype of CHIK was the causative agent of the outbreak in Maceió, Alagoas. This genotype had only been previously identified as circulating in Feira de Santana and Salvador, the two most densely populated regions in Bahia State, in 2014 and 2015 respectively (7, 8). The viral mutation E1-A226V that confers increased transmission in *Aedes albopictus* (9) was not detected in the ECSA-Brazil clade. The most recent common ancestor of the ECSA-Brazilian clade was dated around Jul 2014 (Apr-Aug 2014), in agreement with the date of arrival of the probable index case who travelled from Angola to Feira de Santana (10).

The first CHIKV case in Alagoas was notified in August 2015, over a year after CHIKV was detected in Bahia; the epidemic peak in Alagoas was reached in May 2016, 9 months after its first detection there (Figure). In Feira de Santana, the time-series of CHIKV notified cases suggests a sharp increase in incidence shortly after the virus was first introduced in the municipality in mid-2014. If the ecological conditions and transmission dynamics of CHIKV in Feira de Santana are similar to those of to ZIKV in this city (11), then it is probable that the second wave of CHIKV transmission in 2015 depleted most susceptible individuals, which would explain the thirteen-fold decline in the number of CHIKV cases in 2016.

**Figure.**
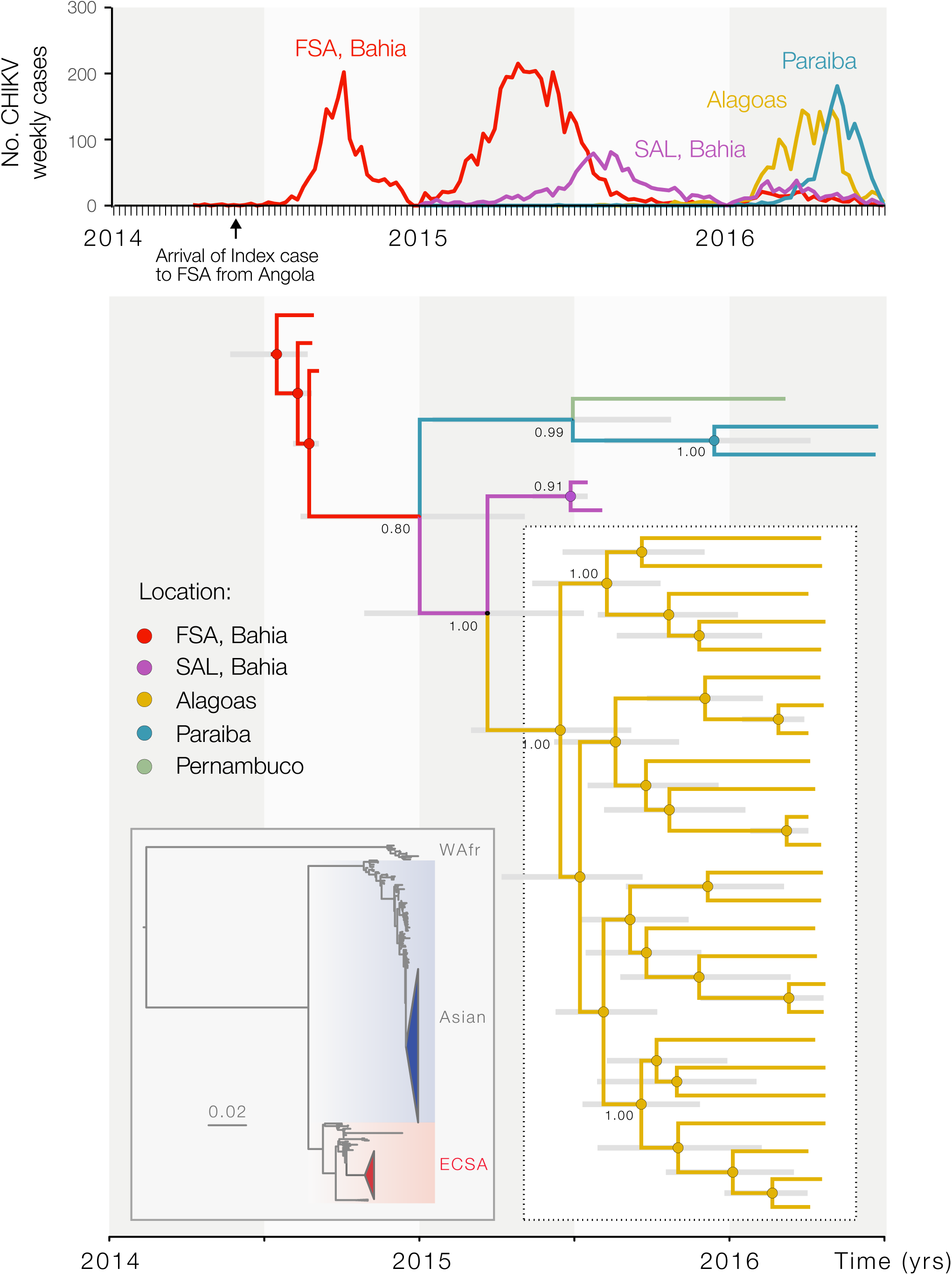
Epidemiological and genetic surveillance of CHIKV in NE-Brazil. The upper panel shows CHIKV notified cases for Alagoas State (Maceió municipality), Paraíba State (João Pessoa municipality), and for Feira de Santana (FSA) and Salvador (SAL) municipalities, both located in Bahia State. The lower right panel shows the molecular clock phylogeny obtained using the 33 novel partial and complete CHIVK sequences (with length > 1,500 nt) collected in the NE Brazil (yellow branches, highlighted in white box). Colours in branches represent most probable locations inferred using a discrete trait diffusion framework (14). At each node, size of the grey circles is proportional to clade location posterior probability. On the left, the inset shows a ML phylogeny with all publicly available CHIKV full-length genomes as of 17 Feb 2017 (*n*=659). WAfr: West African genotype. The IOL genotype has been collapsed. Coloured triangles represent clades circulating in the Americas; the American-ECSA lineage reported in this study is shown in red and the American-Asian lineage is shown in blue.

The novel Maceió CHIKV sequences generated in this study (n=33) fall in a single well supported monophyletic clade (Bootstrap support = 0.89, clade posterior support = 1.00), suggesting that the outbreak was caused by a single founder virus that arrived in Alagoas around Jun 2015 (Jan-Oct 2015); the first CHIKV cases to be notified there were in Aug 2015 (**Figure 1**). It is possible that a second ECSA-CHIVK transmission wave will hit Alagoas and Paraíba in 2017.

Before this study, only 12 Brazilian partial or complete genomes have been reported. Of these, 6 belonged to the Asian genotype, and 6 to the ECSA genotype. Of the former 6, 5 represent imported cases and only one isolate (Accession Number: KP164567) represents an autochthonous case (from Amapá State, which borders French Guiana). Therefore, almost all (98%) of the 40 autochthonous CHIKV cases in Brazil sequenced thus far (33 of which are reported in this study), 98% belong to the ECSA genotype. Further, autochthonous ECSA-CHIKV isolates represent strains sampled in NE-Brazil.

This suggests a pattern of CHIKV infection in NE Brazil that may be distinct to that in the rest of the Americas, where the Asian genotype predominates. NE-Brazil has had the highest number of ZIKV infections and microcephaly cases in Brazil (12). Combined genomic and epidemiological surveillance is needed to study potential interactions between ZIKV, CHIKV, and DENV infection and the risk of severe microcephaly and other illnesses (13).

## Acknowledgments

We are grateful to all participants in this study, and to Luciano Monteiro da Silva and Suzana Santos for support. We are also grateful to Lívia Pinhal and Wanderson Kleber de Oliveira, Brazilian Ministry of Health, for sharing epidemiological data. We also thank Illumina, Inc, Sage Science, Inc, Promega Biotecnologia do Brasil, Ltda, Greiner Bio-One Brasil Produtos Médicos Hospitalares Ltda for the donation of reagents and plastics for this project. This work was supported by FAPESP # 2016/01735-2 and 2012/03417-7. NRF is funded by a Sir Henry Dale Fellowship (Wellcome Trust / Royal Society Grant 204311/Z/16/Z). This research received funding from the ERC under the European Union's Seventh Framework Programme (FP7/2007-2013), grant agreement number 614725-PATHPHYLODYN and the United States Agency for International Development Emerging Pandemic Threats Program-2 PREDICT-2 (Cooperative Agreement No. AID-OAA-A-14-00102).

